# Contrastive Cycle Adversarial Autoencoders for Single-cell Multi-omics Alignment and Integration

**DOI:** 10.1101/2021.12.12.472268

**Authors:** Xuesong Wang, Zhihang Hu, Tingyang Yu, Yixuan Wang, Ruijie Wang, Yumeng Wei, Juan Shu, Jianzhu Ma, Yu Li

## Abstract

We have entered the multi-omics era, and we can measure cells from different aspects. When dealing with such multi-omics data, the first step is to determine the correspondence among different omics. In other words, we should match data from different spaces corresponding to the same object. This problem is particularly challenging in the single-cell multi-omics scenario because such data are very sparse with extremely high dimensions. Secondly, matched single-cell multi-omics data are rare and hard to collect. Furthermore, due to the limitations of the experimental environment, the data are usually highly noisy. To promote the single-cell multi-omics research, we overcome the above challenges, proposing a novel framework to align and integrate single-cell RNA-seq data and single-cell ATAC-seq data. Our approach can efficiently map the above data with high sparsity and noise from different spaces to a low-dimensional manifold in a unified space, making the downstream alignment and integration straightforward. Compared with the other state-of-the-art methods, our method performs better on both simulated and real single-cell data. On the real data, the performance improvement on accuracy over the previous methods is up to 55.7% regarding scRNA-seq and scATAC-seq data integration. Downstream trajectory inference analysis shows that our tool can transfer the labels from scRNA-seq to scATAC-seq with very high accuracy, which indicates our method’s effectiveness.

## INTRODUCTION

Single-cell multi-omics methods promise great opportunities to understand the cellular system more comprehensively. To achieve that, we should obtain multi-omics data at the single-cell level, which is not a trivial task. Although multi-omics profiling approaches for the same set of single cells have become available (1), such as single-cell RNA sequencing (scRNA-seq) and single-cell Assay for Transposase Accessible Chromatin sequencing (scATAC-seq), the experiments on large-scale cells are time-consuming and labor-intensive. Consequently, we have the multi-omics data for a group of cells, but the correspondence between different modalities for a single cell is missing (Figure 1a). More specifically, we want to obtain the high-throughput paired multi-omics data for every single cell, referred to *alignment*. On the other hand, even for data within the same modality, the data distribution of different cell types varies from each other. Besides, the data distribution can be inconsistent because of the subtle differences in measurement processes (2), such as measurement time or equipment used, which is referred to as *batch effects*. They should also be considered when studying the correspondence among different modalities. Considering these, we need to integrate different multi-omics data from the same cell type and batch, which is called *integration*. The above tasks are beneficial and interesting, but challenging, considering the distribution shifting within and across different modalities and the sparsity and high dimension of the single-cell data (3, 4, 5).

**Figure 1.**
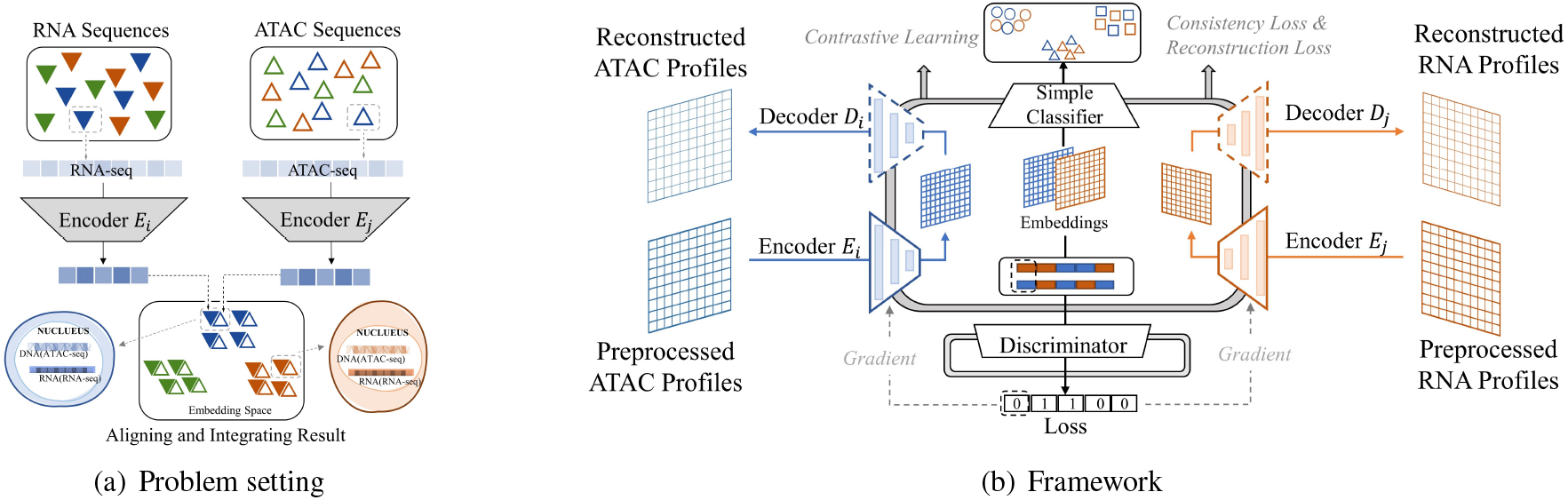
**(a)** scRNA-seq and scATAT-seq data measure different aspects of the same cell. We aim at identifying the correspondence between the two kinds of data from the same set of cells. **(b)** The Con-AAE framework uses two autoencoders to map the two kinds of sequence data into two low dimensional manifolds, forcing the two spaces to be as unified as possible with the adversarial loss and latent cycle-consistency loss. We train the models without pairwise information for the alignment task but consider the data noise explicitly by utilizing self-supervised contrastive learning. We feed the annotated data for the integration task to help the model learn.

Some computational methods have been proposed to deal with these two crucial but challenging problems, aligning and integrating data from different omics. People usually integrate and align multi-omics data in the learned low-dimensional embedding space using dimension reduction techniques, such as Principal Component Analysis (PCA) (6, 7) and nonlinear successors of the classic Canonical Correlation Analysis (CCA) (8). The typical examples are Seurat (9) and Deep Classic Canonical Correlation (DCCA) (10). Seurat(9) relies on the linear mapping of PCA and aligns the embedding vectors based on linear methods Mutual Nearest Neighbors (MNNs) and CCA, which weaken its ability to handle nonlinear geometrical transformations across cellular modalities (11). DCCA can be effective for nonlinear transformation benefiting from deep learning. However, according to the results of our experiments, it is not robust enough when the signal-to-noise ratio (SNR) is low. We also try Maximum Mean Discrepancy (MMD) (12) to replace CCA in the embedding space, but the performance is also not good enough. Several methods requiring no correspondence information are derived under advanced machine learning techniques, such as Pamona (11), MATCHER (13), MMD-MA (14), UnionCom (15), SCOT (16). Although these methods are unsupervised and achieve good performance with encouraging results (16), other additional conditions are still required. For example, MMD-MA and UnionCom, Pamona need the user to specify several hyperparameters, which can be difficult and may need prior information. At the same time, MATCHER and SCOT assume that all datasets share the same underlying structure across cellular modalities (11), which can be ineffective in confronting dataset-specific cell types/structures across the single-cell datasets. Deep learning methods are promising to provide alignment and transfer learning between datasets (17, 18). Deep generative models, such as cycleGAN (19), MAGAN (20), RadialGAN (21) and starGAN (22), are used to learn a nonlinear mapping from one domain to another and achieve great performance on some single modality task. But the above transitions are almost within the same modality and can be disturbed by noise or sparsity in the data (8). The scenario of multi-omics translation and alignment is much more complicated. Some other works propose models to align multi-omics data based on multiple autoencoders (23, 24, 25). However, such methods also can be seriously affected by noise or sparsity, which is a fundamental characteristic of single-cell data.

In general, there are four significant challenges in multi-omics alignment. Firstly, although we have a large amount of unaligned multi-omics data, the aligned data is very scarce. Secondly, the single-cell omics data is of very high dimension and highly sparse. For example, the dropout rate is always extremely high in gene count matrix (26). Furthermore, as we have discussed, the data are highly noisy (3, 4, 5). Finally, although multi-omics data describe the cell status, such as scRNA-seq data and scATAC-seq data, they contain very different information. The mapping between the two kinds of data is highly complicated.

To promote the single-cell multi-omics data analysis, we propose Contrastive Cycle Adversarial Autoencoders (Con-AAE), a framework that can resolve the above challenges precisely (Figure 1b). Con-AAE uses two autoencoders to map the two modality data into two low-dimensional manifolds, forcing the two spaces to be as unified as possible. We use two novel loss terms to achieve that. The first term is called adversarial loss. That is, we combine GAN with autoencoders, forcing the two autoencoders to produce unified embedding to deceive the discriminator, which is designed to distinguish whether two embedding factors are from the same modality. However, only using the adversarial loss may lead to model collapse. To avoid the problem, we further propose a novel latent cycle-consistency loss. For instance, we have two autoencoders for two modalities, ATAC-seq data and RNA-seq data. The embedding produced by the scRNA-seq encoder will go through the scATAC-seq decoder and encoder successively to produce another cycled embedding. We can check the consistency between the original embedding and the cycled embedding. In addition to the above two loss terms, we train the models without pairwise information for the alignment task but consider the data noise explicitly by taking advantage of self-supervised contrastive learning. For the integration task, we train the framework with annotated data. We extensively perform experiments on two real-world datasets, a simulated dataset generated from a real dataset and a group of simulated datasets consisting of various distributions. The two real-world datasets consist of ATAC-seq and RNA-seq data from the same set of cells. The comprehensive experiments on both simulated and real-world datasets show that our method has better performance and is more robust than the other state-of-art methods. The great performance of our method can benefit different kinds of downstream analysis, such as clustering, trajectory inference, pseudotime inference, detection of differential genes, *etc*. Trajectory inference analysis across scRNA-seq and scATAC-seq shows great effectiveness. Although we hide the scATAC-seq information from the inference algorithm, with the help of our method, the label transferred from scRNA-seq data is consistent with the ground-truth label in the scATAC-seq data. Considering the sparsity and difficulty of scATAC-seq data, using information from scRNA-seq data to reach similar performance suggests the effectiveness and the practical usage of Con-AAE.

## MATERIALS AND METHODS

In this section, we give our framework in detail below with Fig. 1b, which illustrates the whole pipeline. To start with, we formalize the alignment problem as,

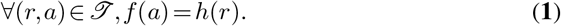

We denote (r, a) as a pair of scRNA-seq and scATAC-seq taken from the same cell. We would like to find two mappings *f* and *h* such that for any aligned {r,a} pairs in *𝒯, f* would map the scRNA-seq profile and scATAC-seq profile to a shared embedding space. Due to the limitations of available real-world aligned data, we are actually working on an unsupervised problem, and the results are evaluated on a few available aligned pairs.

The integration problem could be justified as a clustering problem, and the objective results in finding the corresponding cluster of each scRNA-seq or scATAC-seq profile. The ground-truth labels are available, and, therefore, the model can be trained in a supervised way. For ∀ *x* ∈ {scRNA-seq} ∪ {scATAC-seq}, we want to train a classifier *g* such that

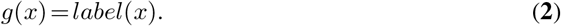

Our primary model is built upon the framework of Adversarial Autoencoders (27) specialized in multi-modality tasks, integrating our novel embedding consistency module and contrastive training process (28). The intuition is that multi-omics from single-cell data should obtain commonality. Their embeddings could live in a unified low-dimensional manifold, making alignment and integration tasks more intuitive.

### Adversarial Autoencoders

The usage of adversarial autoencoders aims to map different omics into a unified latent manifold while able to reconstruct these different aspects. Therefore, as shown in Figure 1b, we are using a coupled set of encoders {*E*_*i*_,*E*_*j*_} (29) to map scATACs-eq,scRNA-seq into manifolds {*Z*_*i*_,*Z*_*j*_}, and decoders {*D*_*i*_,*D*_*j*_} could decode the embedded manifolds back to the original distribution. The reconstruction loss is defined as follows,

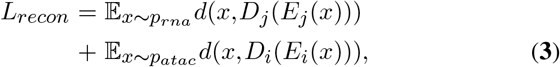

whereas *d* stands for indicated distance in the embedding space. Discriminator *𝒟* tries to align these embedded manifolds and works in the sense that input *x* ∈ *Z*_*j*_, *𝒟* (*x*) = 1 or *x* ∈ *Z*_*i*_, *𝒟* (*x*) = 0.

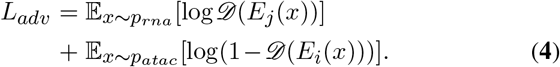

The above losses *L*_*recon*_ and *L*_*adv*_ are trained together with the same weights.

### Latent Cycle-Consistency Loss

The backbone framework enforces the embedding manifolds to align gradually. However, a critical problem underlying is that since scRNA-seq and scATAC-seq data are sparse in a high dimensional domain, the training procedure above only aligns and trains on those regions where the data exist.

For instance, if a region *A* in the embedding space around *E*_*j*_ (*x*^′^),*x*^′^ ∈ {scRNA-seq} does not involve any existing *E*_*i*_(*x*),*x* ∈ {scATAC-seq}. Then, neither the decoder *D*_*i*_ nor the encoder *E*_*i*_ is trained on *A*, thus they would not compute in a “reverse” mapping way, and the result of *E*_*i*_*D*_*i*_*E*_*j*_ (*x*) would be unreasonable or may not lie on the aligned manifold. This critical problem causes the difficulty of inferring from scRNA-seq profile to scATAC-seq profile directly.

Therefore, we introduce a latent consistency loss shown in Figure 2a (19)(30) to resolve this problem,

**Figure 2.**
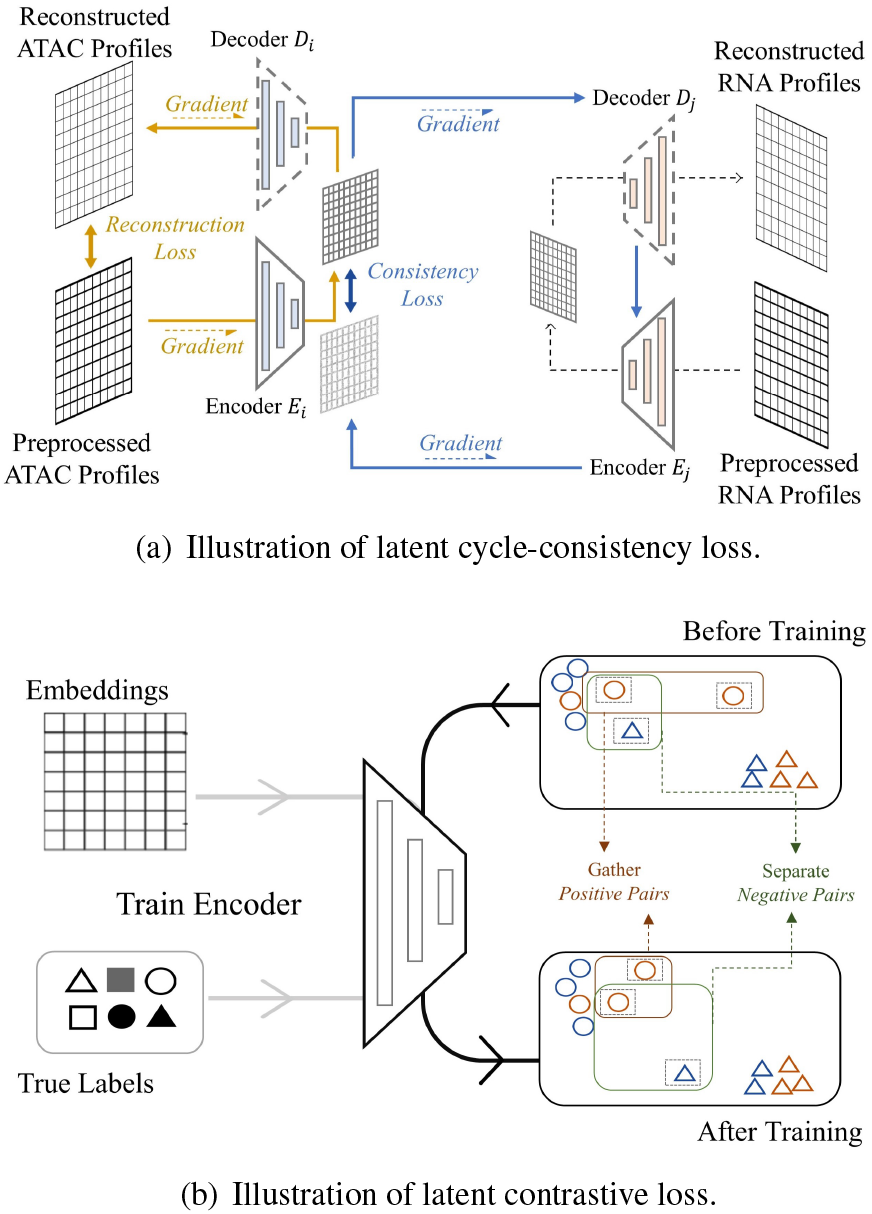
**(a)** The embedding produced by the first encoder will go through the second decoder and encoder successfully to produce another cycled embedding. we can check the consistency between the original embedding and the cycled embedding. **(b)** The contrastive loss minimizes the distance between positive pairs and maximizes the distance between negative pairs. This loss makes our method more robust to noise.

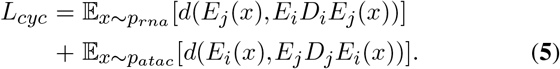

*L*_*cyc*_ aims to train the set of encoder-decoder on the domain where different omics data may not exist, which enforces the smoothness and consistency in those regions. In this way, we could compare the embedding of *E*_*j*_ (*x*),*x* ∈ {scATAC-seq} directly with the existing scRNA-seq embedding around it.

### Supervised Contrastive Loss

The above framework works in an unsupervised manner such that the embedded latent manifolds of multi-omics align properly. A simple classifier trained on our latent space could improve good accuracy of the integration task.

We could further improve our work on both tasks by taking advantage of the ground-truth cell type labels. The cell type labels could refer to biological cell types or the labels of data batches collected from different times or platforms. Following the idea of contrastive learning (31, 32), we employ a contrastive loss in embedding space. It enforces smaller In-Batch distance and larger Between-Batch distance. In-Batch refers to different modality data collected from the same cluster and vice versa. We equally treat both modalities in contrastive training, which benefits the alignment task in the sense that multi-omics of the same single-cell data should obviously belong to the same cluster. We show that lowering the In-Batch distance indeed improves the alignment accuracy in the below ablation studies. On the other hand, contrastive training benefits integration by enabling the decision boundary to be smoother and more robust.

In practice, we first encode data from two modalities to the embedding space. Define the embedding by *z* ∈ ℤ. Given *z*^*a*^ as anchor vector in latent space, we select an *z*^*p*^ such that 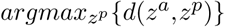, *label* {*z*^*a*^} = *label* {*z*^*p*^}, which is named hard positive. The intuition of hard positive is to find a vector furthest from the anchor within same cluster. Similarly, we have *z*^*n*^ as hard negative such that 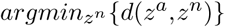, *label* {*z*^*a*^} ≠ *label* {*z*^*p*^}. *z*^*n*^ is defined as the closest vector that from a different cluster. The objective immediately follows,

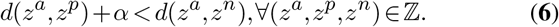

Above, *α* is the margin defined accordingly by us. Thus, by the contrastive loss, we tend to optimize,

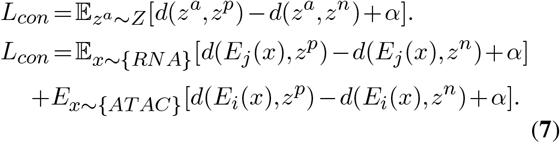

Figure 2b shows that after training, instances within the same cluster are pushed towards each other, and those from the different clusters are forced away. Thus, the decision boundary of the labels tends to be smoother and more robust, which also benefits the alignment task.

### Training Procedure

In the above sections, we proposed several losses related to different objectives. Following the training procedure of Generative Adversarial Nets (23), we adopt a two-stage training scheme where *L*_*adv*_ and *L*_*recon*_,*L*_*cyc*_,*L*_*con*_ are trained separately as the pseudo-code in Algorithm 1.

#### Algorithm 1 Training Procedure

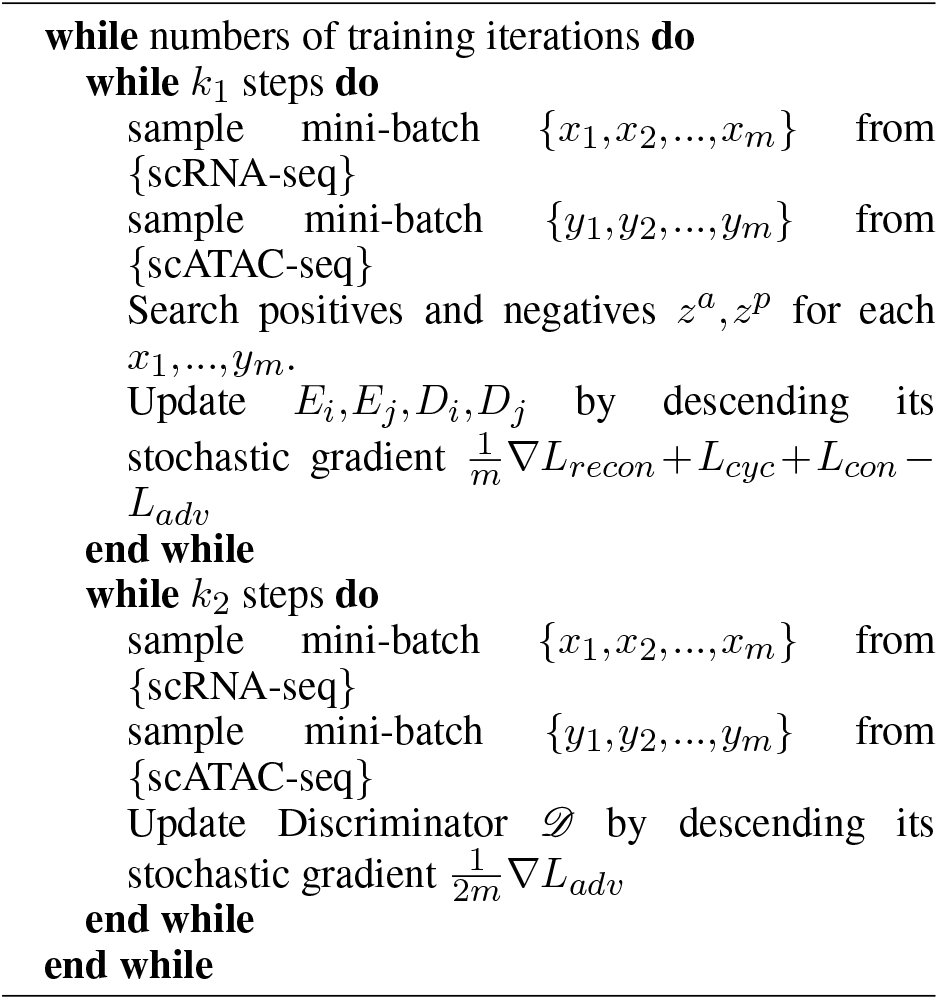

In this way, the Discriminator 𝒟 competes against the encoder-decoder *E*_*i*_,*E*_*j*_,*D*_*i*_,*D*_*j*_ until the training ends and reaches the equilibrium.

## RESULTS

### Experimental Setup

#### Real-world Dataset

We use two sets of single-cell multi-omics data generated by co-assays. The first dataset is generated using the sci-CAR assay (2). For the single-cell ATAC-seq data, we download the processed data from (23), which are computed as described in (2) by counting occurrences of each motif in all accessible sites for each cell, resulting in 815 TF motifs. Then we have a matrix of 1791 × 815 TF motifs. For the single-cell RNA-seq data, we pick the genes with *q* − *value* > 0.05 from the genes being differentially expressed (2), which forms a matrix of 1791 × 2613 genes. Such paired data are collected from human lung adenocarcinoma-derived A549 cells corresponding to 0-, 1-, or 3-hour treatment with Dexamethasone (DEX). So, we have three clusters here, and the cluster label information is available.

We denote the results of SNAREseq (33) assay as the second dataset, which also consists of chromatin accessibility and gene expression. The data are collected from a mixture of human cell lines: BJ, H1, K562, and GM12878. We reduce the dimension of the data by PCA. The resulting matrix for scATAC-seq is of size 1047 × 1000 and 1047 × 500 for gene matrix. The code provided by the author generates annotation information for BJ, H1, K562, and GM12878, so labels of four batches are available.

#### Simulated Datasets

We simulate several datasets of different sizes, which contain 1200, 2100, 3000, and 6000 cells, respectively. For the single-cell RNA-seq data, we utilize three Gaussian distributions with different parameters to generate three batches, and the feature dimension is 1000. For the single-cell ATAC-seq data, we train a four-layer autoencoder with the simulated scRNA-seq data and map them to

500 dimensions as the scATAC-seq data. After that, we randomly set around 40% of features to 0 for RNA-seq data since the real-world scRNA-seq data matrix is usually very sparse. Considering the inevitable mismatch in the experiment, we randomly set around 10% mismatches in the datasets and shuffle all the pairs. Furthermore, we add noise to them with the SNR equal to 5, 10, 15, 20, and 25, along with the version without noise. Then, we have 24 simulated datasets here. Surely, the real-world multi-omics are more complicated than the simulated data, but the experimental results show that our method is sufficient to distinguish the performance of different methods.

In addition to these datasets generated by our method, we also use Splatter(34) to generate a simulated gene count matrix. We call it synthetic RNA-seq, where there are 5000 cells with 1000 genes. Splatter model compute the parameters for the generation with sci-CAR dataset inputted, so we have three clusters in the synthetic RNA-seq data. As described above, we also train a four-layer autoencoder to create the ATAC-seq matrix of 500 dimensions. We call this dataset syn-RNA.

We split all the datasets into training sets and test sets, with the first 80% as the training sets and the last 20% as test sets. Note that we shuffled the data before splitting them.

#### Evaluation criteria

We utilize two existing manners (23) to evaluate integration and alignment, respectively. (a) The fraction of cells whose batch assignment is predicted correctly based on the latent space embedding. (b) *recall*@*k, i*.*e*., the proportion of cells whose true match is within the *k* closest samples in the embedding space (in *ℓ*_1_-distance), while (b) is for alignment.

### Compared with SOTA

Instead of assuming all datasets share the same underlying structure or specifying parts of hyperparameters like some traditional machine learning methods (9, 11, 13, 14, 15, 16), we obtain more information from datasets with partial correspondence information (batch label or cell types label). We select several state-of-art methods based on deep learning like ours, including cross-modal (23), cross-modal-anchor (pairwise information added), DCCA (10), cycle-GAN (19). Moreover, we also compare our method with machine learning methods for integration, including Scanpy (35), Seurat (9), Pamona (11), MMD-MA (14), UnionCom (15), and SCOT (16). Scanpy (35) and Seurat (9) assume that all datasets have the same features, so we apply PCA to make datasets have the same features before using them. We apply Con-AAE and these methods on simulated and real-world datasets.

Firstly, we evaluate all methods on 24 simulated datasets generated by our approach, which are relatively simpler than real datasets and syn-RNA datasets. The extensive experimental results are shown in Figure 4 and 5. As shown in Figure 4, Con-AAE performs better than all the other methods in most cases on the integration task, regardless of data size and SNR. Regarding alignment, we evaluate different methods using *recall*@*k*, whose results are shown in Figure 5. Again, Con-AAE is consistently better than the other competing methods. Notice that our method’s performance is almost consistent against different data sizes and noise levels. In contrast, the other methods may perform well in some settings but poorly in others. The results indicate that Con-AAE is robust and stable enough to have the potential to handle the complicated single-cell multi-omics alignment and integration problems with a low SNR ratio.

Eventually, we care about the methods’ performance on the real-world dataset the most, although the real-world dataset with ground-truth information is limited. We also compare all methods on syn-RNA in this part since syn-RNA is generated from sci-CAR. Still, Con-AAE shows superior performance. On the sci-CAR datasets, Con-AAE outperforms the other methods by up to 36.3% on the integration task, as shown in the upper part of Figure 3. For alignment, Con-AAE always has better performance than all the other methods no matter what *k* is (the bottom part of Figure 3). On the SNAREseq datasets, more obviously, Con-AAE also has dominant performance on each evaluation metric. The improvement on the integration task is up to 55.7% (Figure 3). On the other hand, the performance on *recall*@*k* is better than others no matter what *k* is (Figure 3).

**Figure 3.**
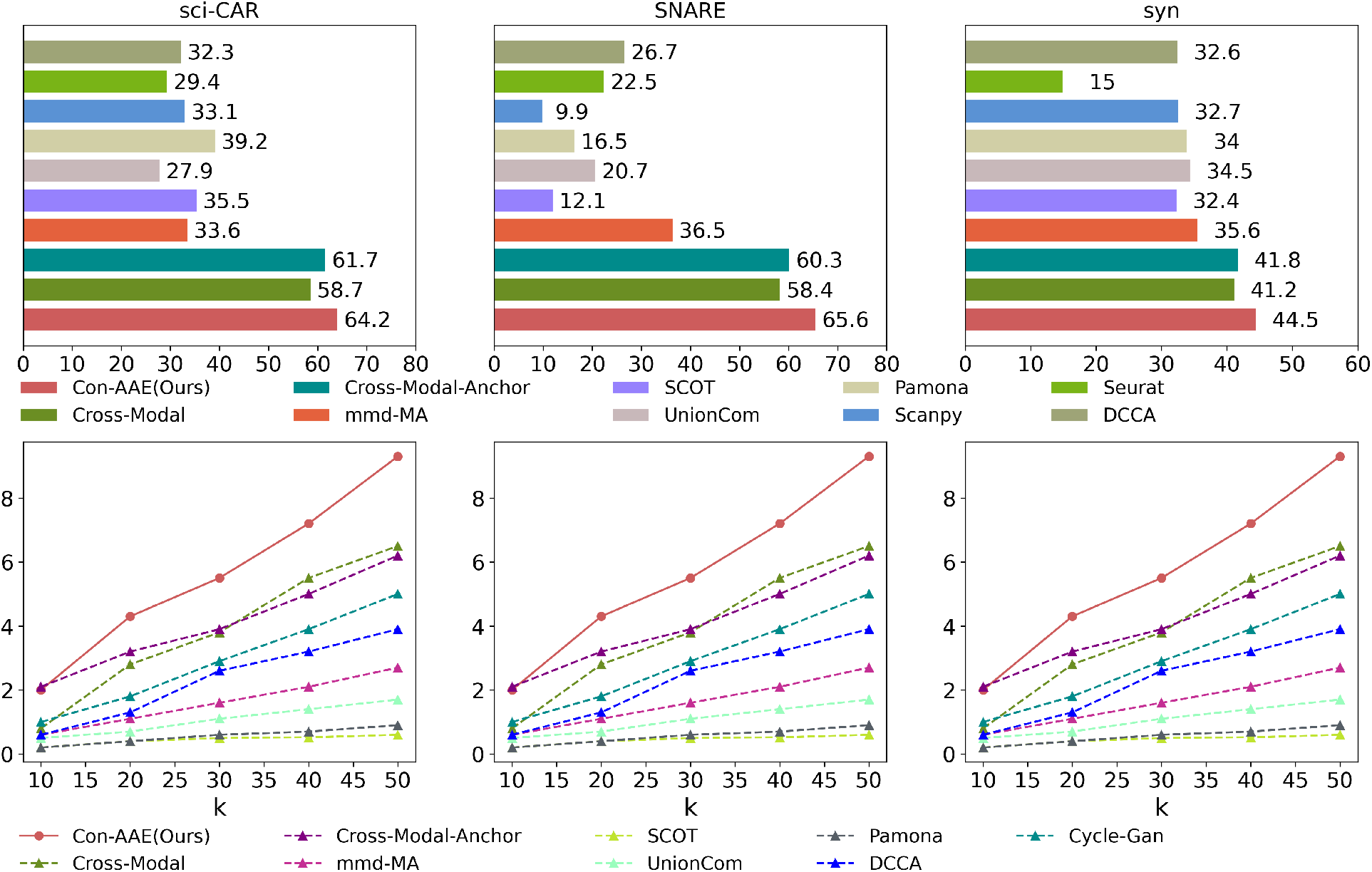
Con-AAE compares with SOTA methods on the two real-world datasets and syn-RNA. The upside is the performance of integration, and the downside is alignment performance. Con-AAE has the best performance on both criteria. Note that the identification of cell pairwise correspondences between single cells is termed “anchor”(9). Cross-modal-anchor indicates that “anchor” information is provided when training Cross-modal.

**Figure 4.**
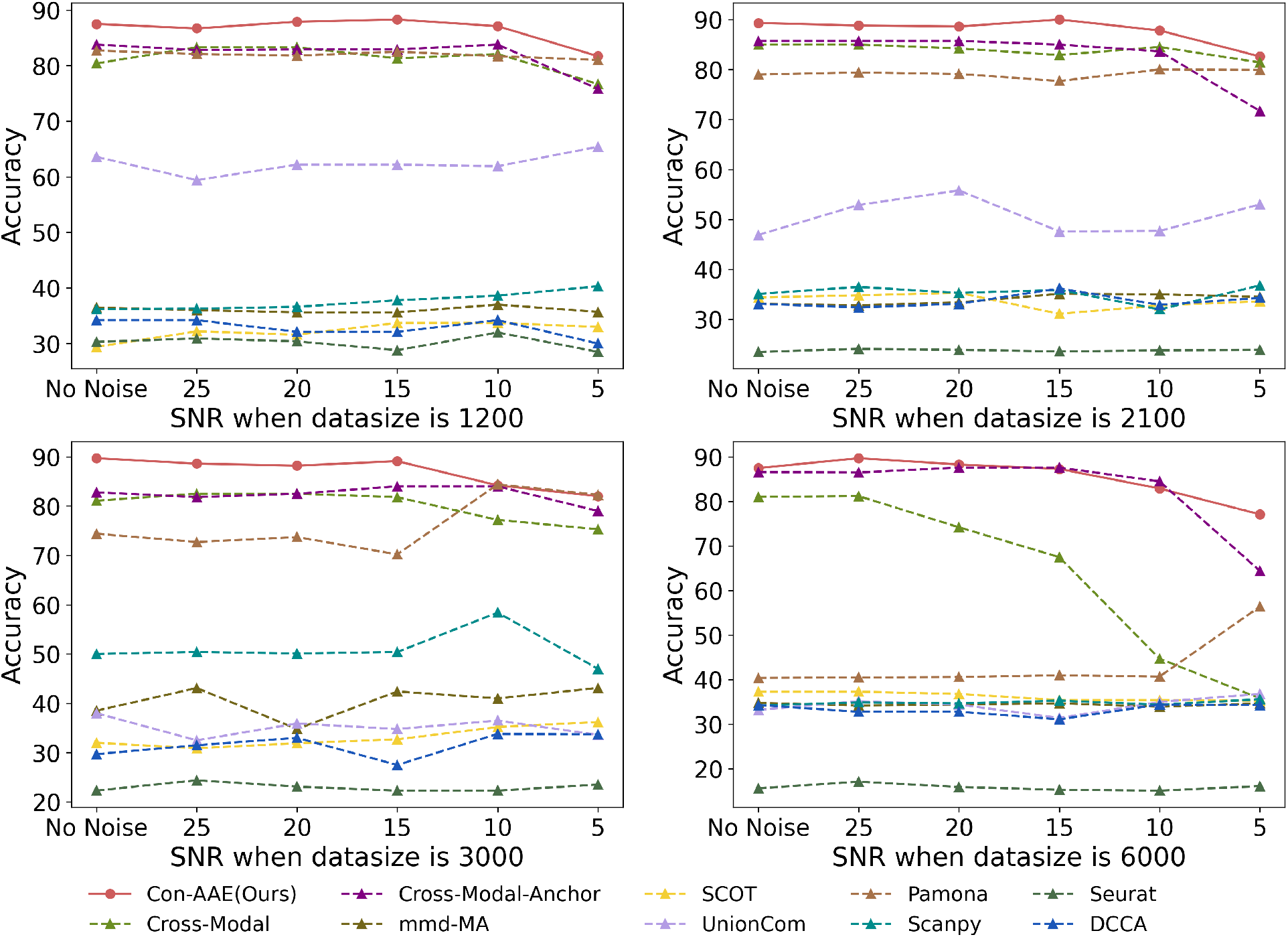
The figure shows the integration performance on 24 simulated datasets with various data sizes and SNR. We can see that Con-AAE almost outperforms other methods all the time. Especially, as the SNR ratio decreases and the size of dataset grows, the performance of all the methods degrades to a different degree. However, Con-AAE still has excellent performance, demonstrating its great scalability and robustness.

**Figure 5.**
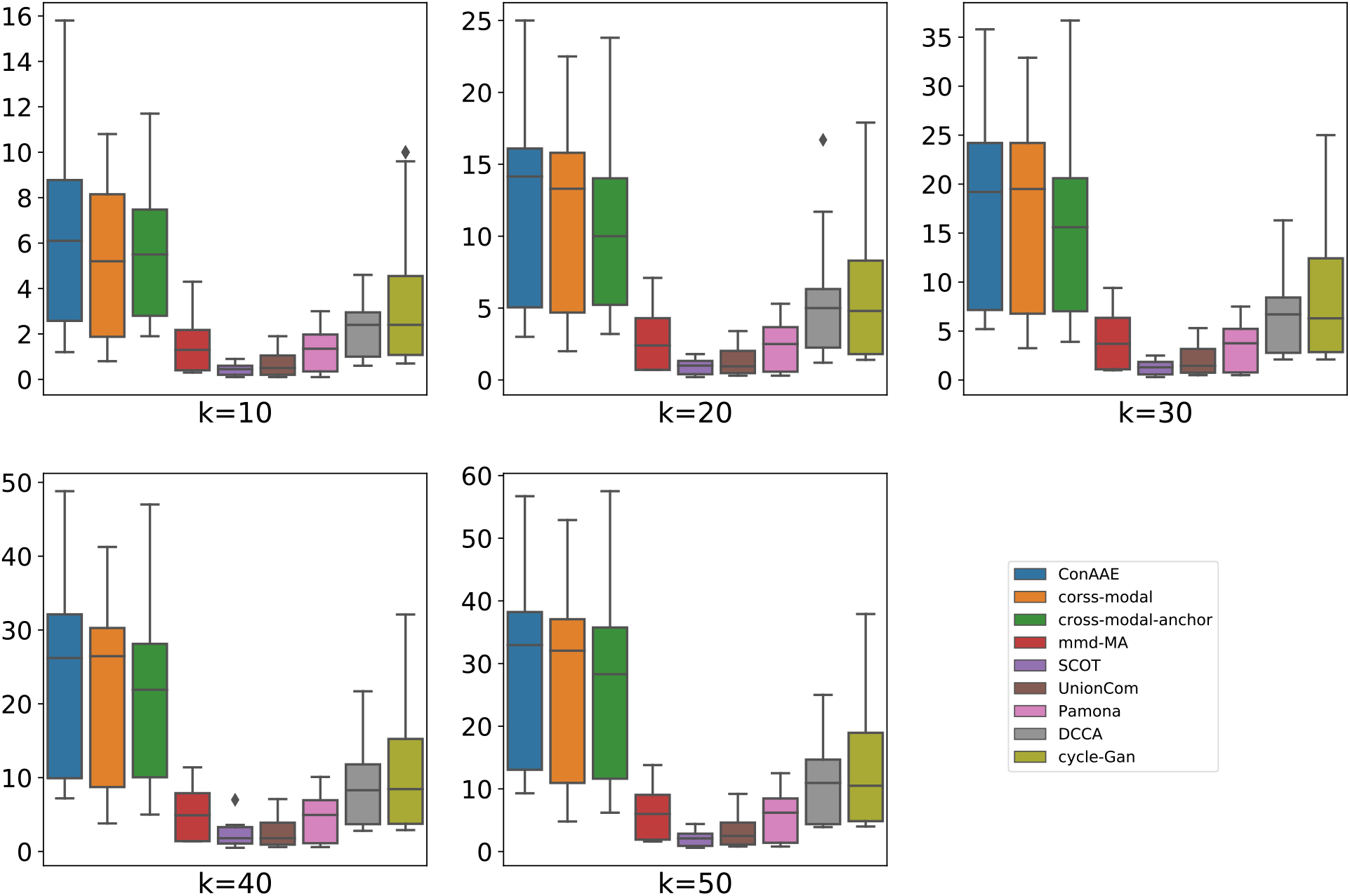
The figure shows the alignment performance on 24 simulated datasets with various data sizes and SNR. Con-AAE almost has the best performance no matter what k is. Corresponding to each K, the box represents the performance on datasets of different sizes and SNR.

### Ablation Studies

We perform comprehensive ablation studies on the sci-CAR dataset, and the results show the effectiveness of different components.

There are three parts in Table 1. The first part indicates there is no adversarial loss in embedding space. The second part indicates an MMD loss (36) instead of an adversarial loss. And the last part indicates whether there is an adversarial loss in the embedding space. Most items in the third part are better than the corresponding items in the other two parts, demonstrating that the adversarial loss works better than MMD loss on this problem.

**Table 1.** Ablation Study of different components in Con-AAE and Comparison with other methods. Basic refers to Coupled AEs plus Simple classifier; adv refers to adversarial loss; mmd refers to mmd loss; anchor refers to pairwise information added; cyc refers to latent cycle-consistency loss; contra refers to contrastive loss.

| Method | Integration<br>ACC | Recall@k<br>k=10 | Recall@k<br>k=20 | Recall@k<br>k=30 | Recall@k<br>k=40 | Recall@k<br>k=50 |
| --- | --- | --- | --- | --- | --- | --- |
| Basic | 56.1 | 3.9 | 9.2 | 13.4 | 18.1 | 20.6 |
| Basic_anchor | 60.8 | 5.3 | 12 | 15.9 | 22.3 | 28.4 |
| Basic_cyc | 56.7 | 6.7 | 11.7 | 15.6 | 19.8 | 24.5 |
| Basic_contra | 58.9 | 6.9 | 12.5 | 15.6 | 21.7 | 27.3 |
| Basic_contra_cyc | 62.2 | 7.8 | 10.8 | 15.9 | 21.2 | 25.9 |
| Basic_mmd | 57.5 | 4.7 | 9.4 | 12.5 | 17.3 | 21.2 |
| Basic_mmd_anchor | 56.4 | 5.3 | 11.4 | 16.7 | 21.2 | 24.3 |
| Basic_mmd_cyc | 60.8 | 3.9 | 10 | 16.2 | 21.2 | 25.1 |
| Basic_mmd_contra | 57.5 | 5.8 | 12.8 | 15.6 | 24 | 27 |
| Basic_mmd_contra_cyc | 57.8 | 4.5 | 9.5 | 13.7 | 19.8 | 24.8 |
| Basic_adv | 58.3 | 4.4 | 10.3 | 14.2 | 17.5 | 23.1 |
| Basic_adv_anchor | 61.7 | 5 | 10.3 | 15.9 | 21.2 | 26.5 |
| Basic_adv_cyc | 58.7 | 5 | 10.3 | 14.2 | 19.8 | 24.3 |
| Basic_adv_contra | 60.61 | 4.4 | 9.4 | 13.6 | 18.4 | 25.4 |
| **Con-AAE** | **63.9** | **5.3** | **12** | **17** | **22.6** | **28.4** |

There are five items in each part of Table 1. The first row represents the basic framework, consisting of two coupled autoencoders and a simple one-layer classifier. The anchor one means pairwise information provided, which indicates that it is a supervised learning model instead of an unsupervised one. “cyc” and “contra” denote latent cycle-consistency loss and contrastive loss, respectively. As shown in the table, adding “cyc” and “contra” improves the model. Apparently, Con-AAE has the best performance. Latent cycle-consistency loss and contrastive loss alone can improve the performance to some degree, but Con-AEE is more robust and has better scalability.

Impressively, Con-AAE has better performance even compared to some supervised methods with the pairwise information provided. Within Table 1, we compare our approach with methods fed pairwise information. We train them using the pairwise information as the supervision for such methods. For Con-AAE, we still perform unsupervised learning using cycle-consistency loss and contrastive loss. Even without the supervised information, Con-AAE can still outperform the basic supervised anchor methods consistently on both tasks. It suggests that cycle-consistency loss and contrastive loss can force our model to learn a unified latent space for the two kinds of single-cell omics data, making the alignment and integration much easier. We also try to combine Con-AAE with the pairwise information. The supervised information can help our method further, but the degree is very slight. We suppose that in the real data, the pairwise information may contain noise, which is common in the single-cell field. Because of the contrastive loss, which makes Con-AAE a robust method, such weak supervision does not help our model too much.

### Visualization

We utilize t-SNE to project the data from the embedding space to the 2D space to visualize the integration and alignment. As we can see in Figure 6 and 7, for the two real-world datasets, we project the scATAC-seq data and scRNA-seq data to a shared embedding space. And within the space, the different omics (with different shapes, such as star and square) data from the same cluster (with the same color) form into clusters, suggesting that our method indeed learned a latent space making it easy to integrate the data from different omics. Furthermore, we use the decoder of each side to translate the embedding vector back to the original space. The resulting data distribution is quite close to the initial distribution, which indicates that our framework learns the underlying features of the data and effectively removes redundant features in encoding.

**Figure 6.**
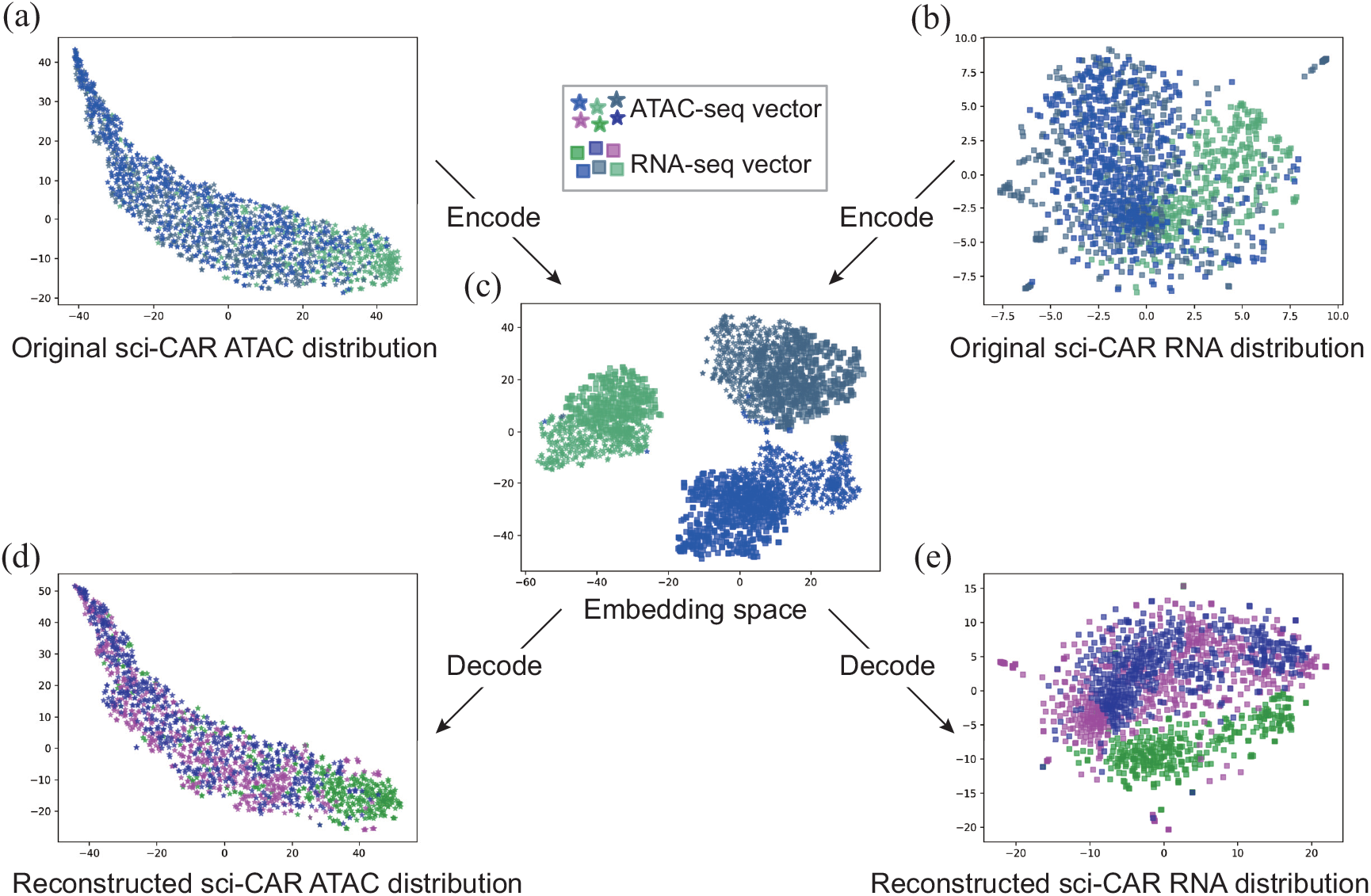
Visualization of the sci-CAR data in the original spaces, shared embedding space, and the reconstructed spaces. The star represents the scATAC-seq data while the square means the scRNA-seq data. Different colors refer to different batch labels, corresponding to 0-, 1-, 3-hours treatment with DEX. For easy viewing, we used different colors when reconstructing data. The distribution of reconstructed data is close to the originality.

**Figure 7.**
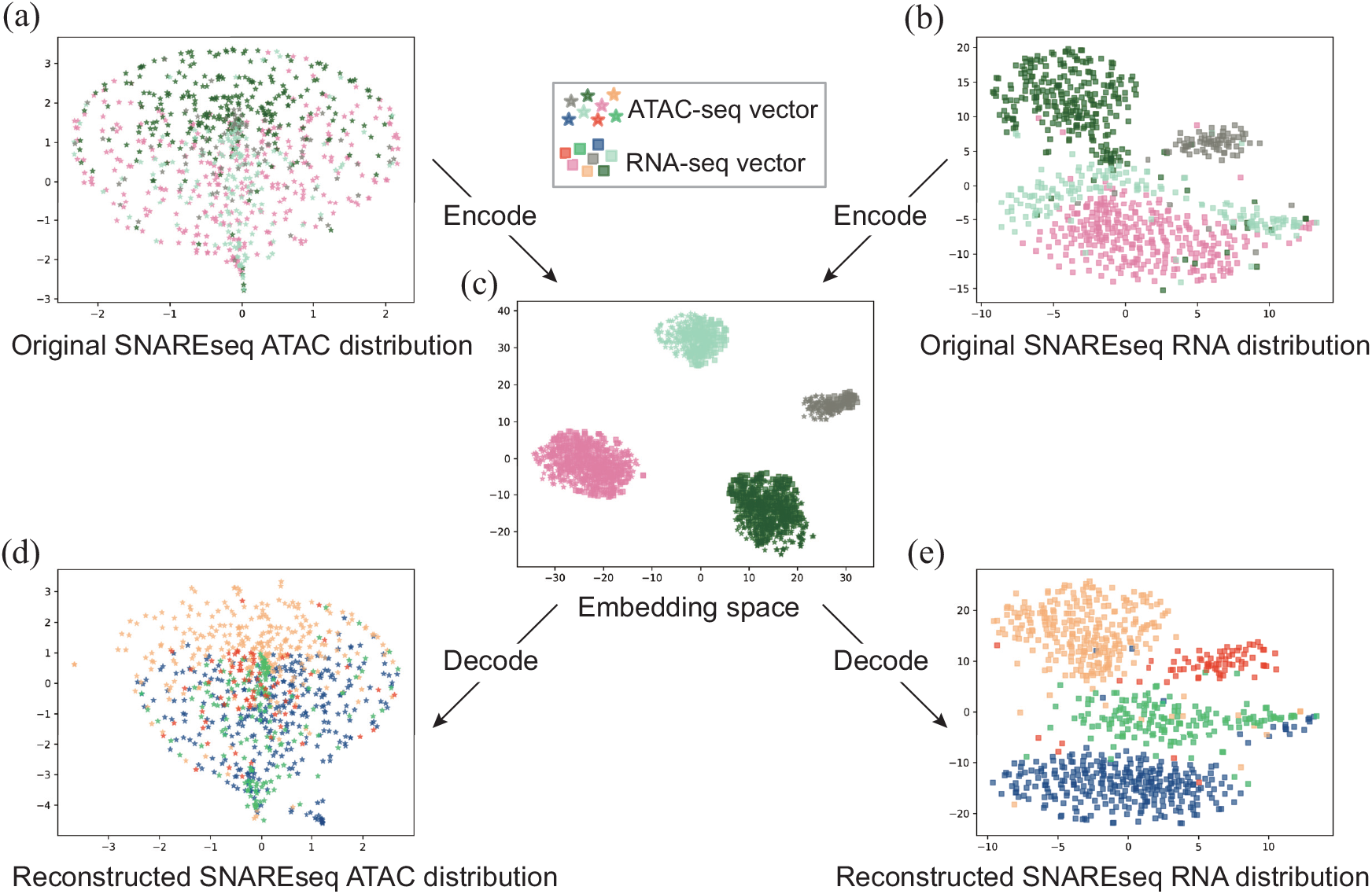
Visualization of the SNAREseq data in the original spaces, shared embedding space, and the reconstructed spaces. The star represents the scATAC-seq data while the square means the scRNA-seq data. Different colors refer to the H1, BJ, K562, and GM12878 human cells. We can see that the scATAC-seq data and scRNA-seq data from the same cell type are tightly combined in the embedding space. And also, we utilize different colors representing reconstructed data, which is quite similar to the original distribution. It further illustrates that our method learns the underlying characteristics of the data.

### Trajectory inference

Suppose that we have one omics data with known cell types, and we want to analyze other omics data from the same population of cells without the label information. For instance, there are lots of work for RNA-seq clustering (9, 37, 38, 39), but little work for ATAC-seq clustering since the sparsity of ATAC-seq poses challenges on computational methods (40). Although obtaining scATAC-seq data from cells is becoming simpler and less expensive (41), getting the clustering information is not an easy task. In this scenario, we can cluster the scRNA-seq data first and then integrate scATAC-seq data with our method to get the cluster information of scATAC-seq data, which can be summarized as label transferring. After that, we can conduct downstream analysis on the scATAC-seq data, such as inferring trajectory, mapping nucleosomes, detecting transcription factor binding sites or differential chromatin states (41). Here, we demonstrate our model on the test set of sci-CAR, which has 359 cells. We integrate scATAC-seq data with scRNA-seq data and transfer the labels from scRNA-seq. We implement trajectory inference on these scATAC-seq data with transferred labels by PAGA (42) and compare it with the trajectory inference derived from original ground-truth labels as shown in Figure 8. Clearly, the trajectory inference results of transferred and original labels are similar, demonstrating our method’s effectiveness and practical usage. In addition, we show that the accessibility of transcription factor motifs changes with the treatment time of DEX, which can help us understand how cells respond to drugs.

**Figure 8.**
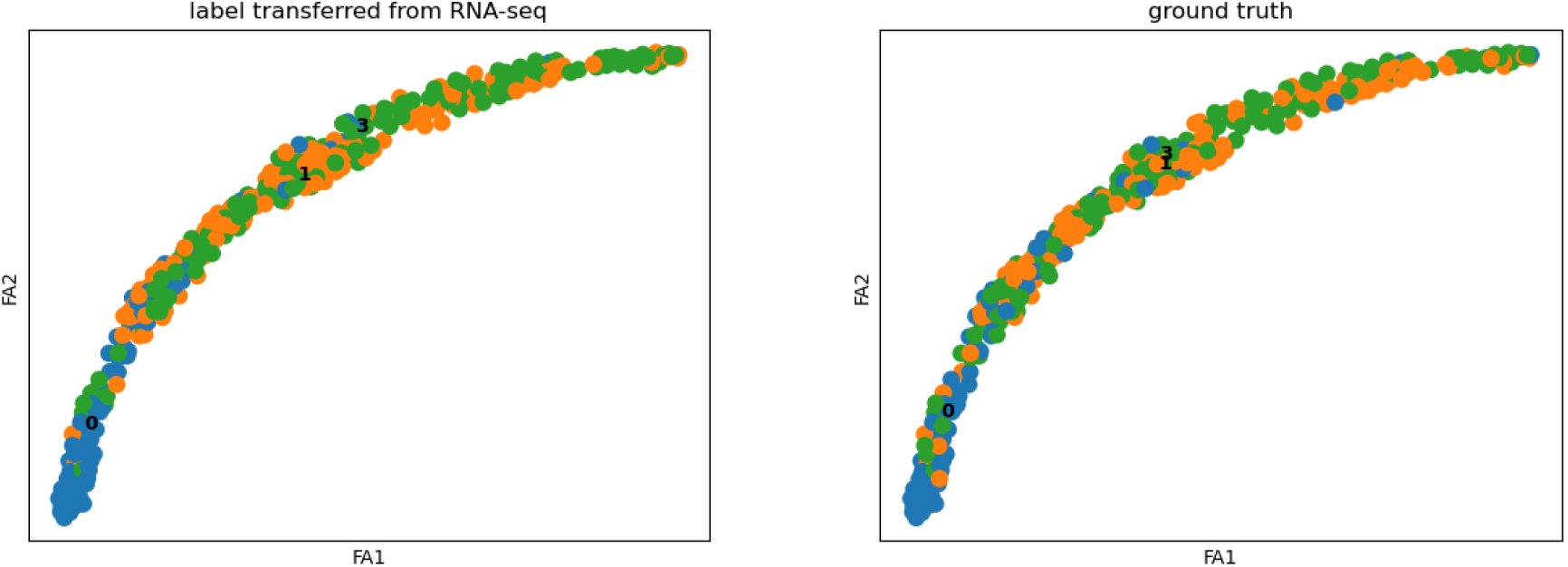
The trajectory inference results of the transferred and original labels are similar. 0,1,3 correspond to 0-, 1-, or 3-hour treatment with DEX. The trajectory inference infers the pattern of cells in the dynamically developmental progress with the treatment of dexamethasone (DEX). The result of labels transferred from scRNA-seq data is very similar to the effects of ground-truth labels, which suggests that our model performs significantly on the integration task.

## DISCUSSION

In this paper, we propose a novel framework, Con-AAE, aiming at integrating and aligning the multi-omics data at the single-cell level. On the one hand, our proposed method can map different modalities into the embedding spaces and overlap these two distributions with the help of an adversarial loss and a novel latent cycle-consistency loss. On the other hand, we apply a novel self-supervised contrastive loss in the embedding space to improve the robustness and scalability of the entire framework. Comprehensive experimental results on the simulated and real datasets show that the proposed framework can outperform the other state-of-the-art methods for both alignment and integration tasks. Detailed ablation studies also dissect and demonstrate the effectiveness of each component in the framework. Trajectory inference on multi-omics data further demonstrates our method’s effectiveness and practical usage. Our method will be helpful for both the single-cell multi-omics research and the general multi-modality learning tasks in computational biology. For future work, we aim to extend our work from a two-domain task to a multiple-domain study, allowing it to integrate and align multiple omics. Besides integration and alignment between sequence modalities, we intend to perform our method on different kinds of biological data, including but not limited to images, geometrical spatial structure, *etc*. Obviously, it is exciting to investigate the spatial transcriptomics data. We will also develop methods for translating modalities. By doing so, we hope to build a system that could benefit various downstream analyses in single-cell multi-omics and spatial multi-omics.

## ACKNOWLEDGEMENTS

This work was supported by a grant by The CUHK Shenzhen Research Institute.

## REPRODUCIBILITY

### Details of the experiments

The details of the network architecture are summarized in Table 1. The learning rate of the model is 0.0001, and the batch size is 32 for the real-world dataset, 100 for simulated datasets. We train this model for 4000 epochs using the Adam optimizer with *β*_1_ = 0.5, *β*_2_ = 0.999, and weight decay is set to 0.0001. Using early stopping, we find that the best model usually appears in 1000 epoch or 2000 epoch. The activation function is LeakyReLU after each layer, followed by Batch Normalization.

**Table 1.** “d” refers to the dimension of the input data. 50 is the dimension of the embedding space. Note that the last hidden layer in the encoder and the fast layer in the decoder are 100.

|  | Encoder | Decoder |
| --- | --- | --- |
| Input size | d | 50 |
| Hidden layer size | d, d, d, 100 | 100, d, d, d |
| Output size | 50 | d |

The discriminator consists of 2 hidden layers with 50 and 100 nodes, respectively. The output size is 1. The simple classifier consists of 1 layer, and the output size is 3.

### Implementation of Con-AAE

We have implemented the Con-AAE in Python 3.7.7 with Pytorch 1.0, whose experiments were run on Nvidia Tesla P100. The source code is available at https://github.com/kakarotcq/RNA-Seq-and-ATAC-Seq-mapping.

### The hyperparameter tunning range

For all the methods, the weights of different loss terms are shown in Table 2. The weights of anchor loss and cycle-consistency loss are supposed to be low because the noise level of the real dataset is high, with a few mismatches, while contrastive loss should be adjusted to a higher value.

**Table 2.** Loss terms and their corresponding weights for training Con-AAE.

| Loss function | Loss type | Weight |
| --- | --- | --- |
| Reconstruction loss | MSELoss | 10.0 |
| Adversarial loss | MSELoss | 10.0 |
| Classifier loss | CrossEntropyLoss | 1.0 |
| Cycle-consistency loss | MSELoss | 1.0, 5.0, 10.0 |
| Contrastive loss | MarginRankingLoss | 1.0, 5.0, 10.0 |
| Anchor loss | MSELoss | 0.05, 0.1, 1.0 |

## EXTRA EXPERIMENTAL RESULTS

We also try to train Con-AAE with the pairwise information, denoted as Con-AAE-anchor. The comparison between Con-AAE and Con-AAE-anchor is shown in Table 3. We can see that the integration performance is almost the same while the alignment performance is improved slightly as the pairwise information is added. In general, Con-AAE is a very stable unsupervised method, which is robust to noise. So the weak supervised pairwise information with a high level of noise does not significantly improve Con-AAE’s performance. But still, this experiment suggests that Con-AAE is a flexible framework that can incorporate supervised information.

**Table 3.** Comparison between Con-AAE and Con-AAE-anchor on sci-CAR dataset. Con-AAE is a very stable unsupervised method, which is robust to noise. So the weak supervision with a high level of noise does not significantly improve Con-AAE’s performance.

| Method | Integration ACC | Recall@k=10 | Recall@k=20 | Recall@k=30 | Recall@k=40 | Recall@k=50 |
| --- | --- | --- | --- | --- | --- | --- |
| Con-AAE | 63.9 | 5.3 | 12 | 17 | 22.6 | 28.4 |
| Con-AAE-anchor | 63.4 | 7 | 13.7 | 18.4 | 25.4 | 29.1 |

We also conduct ablation studies on the simulated datasets. The quantitative results are shown in the Table 4 and Figure 1. We can see that the cycle-consistency loss and contrastive loss could improve the model in many cases, but not stably enough. Compared with that, Con-AAE almost always has the best performance with the change of data sizes and SNRs, while the other methods struggle when the SNRs or data sizes change.

**Table 4.** Ablation study on the simulated datasets with various SNRs: basic plus adversarial loss (denoted as Basic adv), basic plus adversarial loss and cycle-consistency loss method (denoted as Basic adv cyc), basic plus adversarial loss and contrastive loss (denoted as Basic adv contra) and basic plus adversarial loss, cycle-consistency loss, and contrastive loss (Con-AAE). The performance of Con-AAE is in bold. Clearly, Con-AAE’s performance is very stable across different data sizes and SNRs, compared to the other baseline methods.

| #Sample | Method | No Noise | SNR25 | SNR20 | SNR15 | SNR10 | SNR5 | SD | AVG |
| --- | --- | --- | --- | --- | --- | --- | --- | --- | --- |
| 1200 | Basic_adv | 80.4 | 83.3 | 83.3 | 81.3 | 82.1 | 76.7 | 2.47 | 81.18 |
|  | Basic_adv_cyc | 83.3 | 83.3 | 83.8 | 83.3 | 84.2 | 82.3 | 0.64 | 83.36 |
|  | Basic_adv_contra | 85.0 | 85.4 | 85.0 | 87.5 | 83.8 | 72.5 | 5.38 | 83.20 |
|  | **Con-AAE** | **87.5** | **86.7** | **87.9** | **88.3** | **87.1** | **81.7** | **2.43** | **86.53** |
| 2100 | Basic_adv | 85.0 | 85.0 | 84.2 | 82.9 | 84.5 | 81.4 | 1.42 | 83.80 |
|  | Basic_adv_cyc | 72.4 | 72.4 | 71.2 | 70.4 | 76.2 | 77.4 | 2.82 | 73.33 |
|  | Basic_adv_contra | 91.0 | 90.0 | 91.9 | 91.4 | 86.2 | 72.1 | 7.63 | 87.10 |
|  | **Con-AAE** | **89.3** | **88.8** | **88.6** | **90.0** | **87.8** | **82.6** | **2.67** | **87.85** |
| 3000 | Basic_adv | 81.1 | 82.5 | 82.5 | 81.8 | 77.2 | 75.3 | 3.06 | 80.07 |
|  | Basic_adv_cyc | 85.8 | 86.8 | 87.0 | 86.5 | 84.0 | 80.6 | 2.47 | 85.12 |
|  | Basic_adv_contra | 88.8 | 90.0 | 89.2 | 67.5 | 60.8 | 86.5 | 12.87 | 80.47 |
|  | **Con-AAE** | **89.7** | **88.6** | **88.2** | **89.1** | **74.2** | **82.0** | **6.12** | **85.30** |
| 6000 | Basic_adv | 81.0 | 81.2 | 74.2 | 67.5 | 44.7 | 35.8 | 19.33 | 64.07 |
|  | Basic_adv_cyc | 81.1 | 85.2 | 85.5 | 78.6 | 79.2 | 78.5 | 3.24 | 81.35 |
|  | Basic_adv_contra | 90.8 | 90.9 | 91.3 | 90.0 | 81.9 | 67.6 | 9.43 | 85.42 |
|  | **Con-AAE** | **87.5** | **89.7** | **88.3** | **87.3** | **82.9** | **77.1** | **4.69** | **85.47** |

**Figure 1.**
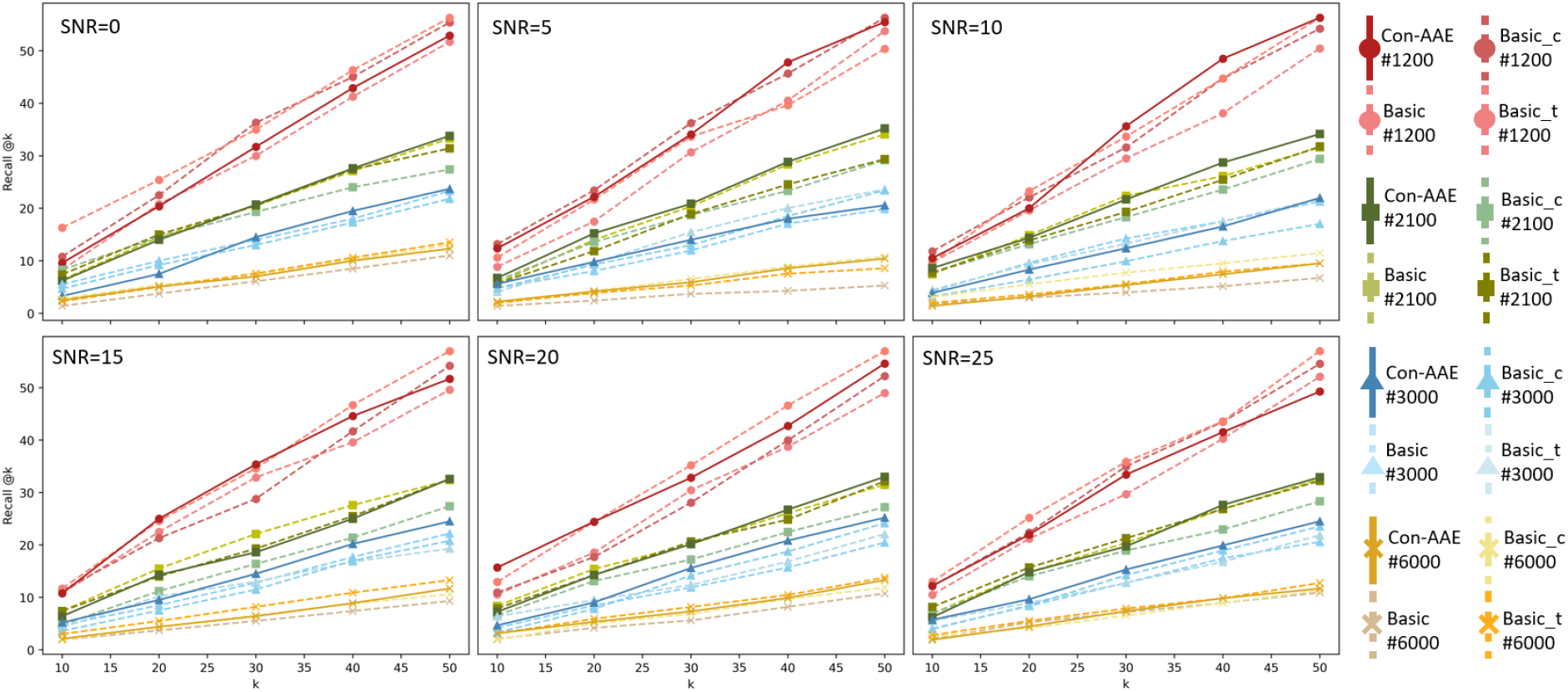
The alignment performance on simulated datasets with different components.

